# Differential neural circuitry behind autism subtypes with imbalanced social-communicative and restricted repetitive behavior symptoms

**DOI:** 10.1101/2020.05.08.083758

**Authors:** Natasha Bertelsen, Isotta Landi, Richard A. I. Bethlehem, Jakob Seidlitz, Elena Maria Busuoli, Veronica Mandelli, Eleonora Satta, Stavros Trakoshis, Bonnie Auyeung, Prantik Kundu, Eva Loth, Guillaume Dumas, Sarah Baumeister, Christian F. Beckmann, Sven Bölte, Thomas Bourgeron, Tony Charman, Sarah Durston, Christine Ecker, Rosemary J. Holt, Mark H. Johnson, Emily J. H. Jones, Luke Mason, Andreas Meyer-Lindenberg, Carolin Moessnang, Marianne Oldehinkel, Antonio Persico, Julian Tillmann, Steven C. R. Williams, Will Spooren, Declan G. M. Murphy, Jan K. Buitelaar, the EU-AIMS LEAP group, Simon Baron-Cohen, Meng-Chuan Lai, Michael V. Lombardo

## Abstract

Social-communication (SC) and restricted repetitive behaviors (RRB) are autism diagnostic symptom domains. SC and RRB severity can markedly differ within and between individuals and may be underpinned by different neural circuitry and genetic mechanisms. Modeling SC-RRB balance could help identify how neural circuitry and genetic mechanisms map onto such phenotypic heterogeneity. Here we developed a phenotypic stratification model that makes highly accurate (97-99%) out-of-sample SC=RRB, SC>RRB, and RRB>SC subtype predictions. Applying this model to resting state fMRI data from the EU-AIMS LEAP dataset (n=509), we find that while the phenotypic subtypes share many commonalities in terms of intrinsic functional connectivity, they also show replicable differences within some networks compared to a typically-developing group (TD). Specifically, the somatomotor network is hypoconnected with perisylvian circuitry in SC>RRB and visual association circuitry in SC=RRB. The SC=RRB subtype show hyperconnectivity between medial motor and anterior salience circuitry. Genes that are highly expressed within these networks show a differential enrichment pattern with known autism-associated genes, indicating that such circuits are affected by differing autism-associated genomic mechanisms. These results suggest that SC-RRB imbalance subtypes share many commonalities, but also express subtle differences in functional neural circuitry and the genomic underpinnings behind such circuitry.

---

Autism spectrum disorder (ASD) is a clinical consensus label used to characterize individuals with a collection of early onset developmental difficulties in the domains of social-communication (SC) and restricted repetitive behaviors (RRB)^1,2^. The single diagnostic label of autism helps many individuals in a variety of ways by being incorporated into a sense of identity, explaining challenging aspects of life, and/or enabling access to services. However, the diagnosis also encapsulates a vast amount of multi-scale heterogeneity. In the face of such heterogeneity, future translational research must develop a deeper understanding of how biological mechanisms affect individuals and must develop more personalized approaches towards interventions to help facilitate positive outcomes^3^.

Because heterogeneity manifests across every scale from phenome to genome, it is important to understand whether top-down phenotypic stratifications may be useful. For example, there is evidence to suggest that important phenotypic stratifications could be made based on the balance between SC and RRB domains. Prior work has suggested that SC and RRB domains could be fractionable at behavioral^4^ and neural levels^5–7^ and potentially underpinned by different genetic mechanisms^8–11^. However, robust evidence of this phenotypic fractionation mapping onto differential neural circuitry and genomic mechanisms has yet to be identified. The potential multiscale fractionation of these domains provides a strong starting point for understanding how multiscale heterogeneity manifests in autism from genome to phenome.

In this work, we test the hypothesis that subtyping individuals by the degree of SC-RRB balance can help identify differential biological mechanisms. Past research utilizing ‘gold standard’ diagnostic tools such as the Autism Diagnostic Observation Schedule (ADOS) and the Autism Diagnostic Interview-Revised (ADI-R) (e.g., ^12–14^) have suggested the presence of 3 SC-RRB balance subtypes: 1) medium to high levels of both SC and RRB severity (SC=RRB); 2) medium to high SC severity and comparatively lower RRB severity (SC>RRB); and 3) medium to high RRB severity and comparatively lower SC severity (RRB>SC). These subtypes might be underpinned by a common pathway if they showed similar neural circuits and genomic mechanisms that differ from a typically-developing (TD) comparison group. However, based on the hypothesis that SC and RRB domains are fractionable across multiple levels, it could be that these subtypes diverge onto multiple atypical pathways from genome to phenome^15^ (Fig. 1). This idea has not yet been tested with respect to macroscale neural circuitry and its link to functional genomic mechanisms. Here we evaluate how SC-RRB balance subtypes link up to differential macroscale connectome phenotypes, measured with resting state fMRI (rsfMRI) functional connectivity. Functional connectivity networks are known to be linked to underlying transcriptomic mechanisms, particularly with regards to the spatial patterning of gene expression across the brain (e.g., ^16–18^). Given that subtypes could exhibit different functional connectome phenotypes, we leverage the link between macroscale rsfMRI functional networks and transcriptomic mechanisms to better understand whether autism-relevant functional genomic mechanisms differentially affect such phenotypes.

**Fig. 1:**
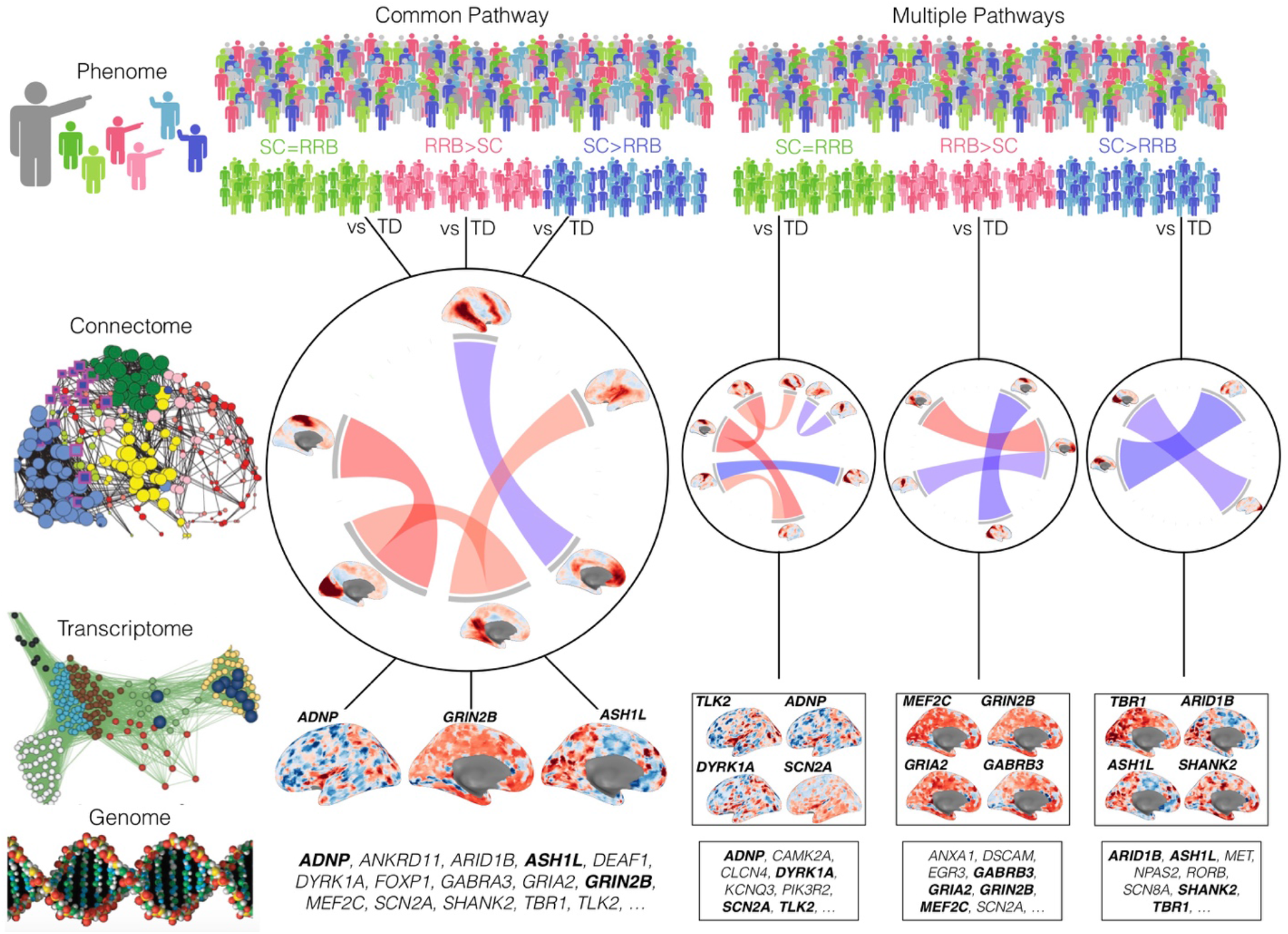
Approach towards testing common pathway versus multiple pathways explanations behind SC-RRB balance in autism. In this figure we depict two alternatives for how SC-RRB balance subtypes (phenome level; SC=RRB, green; RRB>SC, pink; SC>RRB, blue) could be explained at the level of macroscale functional connectome phenotypes measured with rsfMRI (connectome level) and autism-associated functional genomic mechanisms (e.g., transcriptome and genome levels). Columns in this figure depict the common pathway (middle) and multiple pathways (right) models. The common pathway model predicts that when each subtype is compared to a typically-developing (TD) comparison group, they converge and share a common difference from TD in affected macroscale rsfMRI functional connectome phenotype. Underpinning this shared connectome phenotype are a myriad of differing functional genomic mechanisms. At the level of the transcriptome, we identify genes linked to macroscale functional networks by identifying genes whose spatial expression pattern across the brain is similar to the spatial topology of the macroscale functional network. This procedure generates a list of genes relevant for such macroscale networks and these lists are then tested for enrichment in autism-association functional genomic mechanisms. The gene list at the genome level represents an example of possible autism-associated genes that may (bold) or may not (non-bold) be linked to macroscale functional networks. In contrast to the common pathway model, the multiple pathways model would highlight that differential connectome phenotypes when compared to TD are unique to each subtype, and that each of these subtype-specific connectome phenotypes is underpinned by a differing set of autism-associated functional genomic mechanisms.

To test these ideas, we developed a stratification approach that subtypes individuals based on the within-individual balance between SC versus RRB severity, as measured by ADI-R data from thousands of individuals (n=2,628) obtained from the National Database for Autism Research (NDAR) (https://nda.nih.gov). This approach can be used to make highly accurate out-of-sample subtype predictions and thus can be applicable to any new dataset where ADI-R data is collected. We applied this SC-RRB balance stratification model to the deeply phenotyped EU-AIMS LEAP dataset^19–21^ to examine how functional connectivity may differ between SC-RRB balance subtypes or relative to a typically-developing (TD) comparison group. While SC-RRB balance subtypes are important to test, it may be also useful to consider overall level of SC and RRB severity as important and to characterize this severity in a continuous manner. Thus, we also evaluate other possible continuous/dimensional models that examine SC and RRB separately as well as using SC-RRB balance score as a continuous variable. Finally, in order to link functional connectome phenotypes to autism-associated genes, we utilize the Allen Institute Human Brain Atlas^22,23^ to identify genes whose spatial expression pattern is highly similar to macroscale functional networks that differ amongst the SC-RRB subtypes. These functional network-relevant gene lists are then investigated for enrichment in a variety of autism-associated gene lists derived from evidence at genetic or transcriptomic levels. This will allow for tests of the hypothesis that subtype disruption of imaging-derived phenotypes preferentially occurs to macroscale networks with high levels of gene expression of autism-associated genes^24^. This approach will also allow us to test whether autism-associated genes affect networks similarly or differently across the SC-RRB balance subtypes.

## Materials and Methods

### NDAR Datasets

For the initial set of analyses to derive the approach to characterizing SC and RRB symptom domains, we utilized ADI-R from the National Database for Autism Research (NDAR). Because individuals can differ over the lifespan with regard to current symptom presentation, we opted for using the ADI-R (‘diagnostic algorithm’ scores) as the measure of symptomatology because it allows for assessment of symptoms at similar ages regardless of the age of the participant at the time of testing. This is an important point within the context of the current study, whereby the age range of the follow-up sample (i.e. EU-AIMS LEAP) is notably wide (i.e. 6-30 years). Within the SC domain, many items are rated based on presentation around 4-5 years of age. This is required because such items probe behaviors that are developmentally most appropriate in typical development around this point in the lifespan and the absence of such behaviors in autistic individuals is of diagnostic importance. In contrast, for many types of RRB domain items, the presence rather than the absence of such behaviors is of diagnostic importance for autism. For these items, the behaviors could occur at any point during the lifespan and are not developmentally restricted to a specific age range.

On December 13, 2019 we conducted a search of NDAR to extract all datasets utilizing the ADI-R^25^. This resulted in 60 independent datasets totaling 2,628 unique individuals. From here, we filtered for all individuals who had data for the verbal communication items (e.g., acquisition of words, phrases, social verbalization, chit-chat, reciprocal conversation), leaving a total of 1,781 individuals across 57 independent datasets. Within each of these 57 datasets, we randomly split the dataset in half to achieve independent Discovery and Replication sets that are balanced across the 57 datasets and by sex (Discovery n=889, mean age = 8.91 years, SD age = 5.26 years, 77% male; Replication n=890, mean age = 8.89 years, SD age = 5.37 years, 77% male). See Table 1 for characteristics of the NDAR Discovery and Replication sets. ADI-R item-level data was used to rescore algorithm totals according to the DSM-5^26^ symptom dyad of social-communication (SC) and restricted repetitive behavior (RRB) domains. SC is comprised of 3 subscales (A1, A2, A3), while RRB is comprised of 4 subscales (B1, B2, B3, B4). See Supplementary Table 1 for how items break down into each domain and subscale within a domain. Only item scores of 0 to 3 (indicating increasing SC or RRB symptom severity) were utilized, while scores of 6 to 9 (dummy scores, not indicating symptom severity) were not used. Scores of 3 were kept as is (i.e. not converted to 2 as would typically occur when scoring the ADI-R algorithm) in order to retain information about severity conveyed by the difference between a score of 2 versus 3. Because the number of items in each subscale can vary depending on a person’s age (see Supplementary Table 1) and by the number of items with possible scores of 0 to 3, we used percentage scores in order to ensure that the estimates of severity on each subscale are on a comparable scale across individuals. These percentage scores for each domain subscale were then summed and scaled by number of subscales to achieve the final domain total percentage severity.

**Table 1:** Participant characteristics from the NDAR datasets. At a z-threshold of 1, this table shows sample sizes and descriptive statistics (mean and standard deviation) for age and ADOS social affect (SA) and restricted repetitive behaviors (RRB) calibrated severity scores. The final column on the right shows the F-statistic and p-value from an ANOVA testing for an effect of group. ^1^ Sample sizes: ADOS (Discovery, RRB>SC n=19, SC=RRB n=99, SC>RRB n=35; Replication RRB>SC n=8, SC=RRB n=119, SC>RRB n=26); FIQ (Discovery, RRB>SC n=39, SC=RRB n=142, SC>RRB n=18; Replication RRB>SC n=40, SC=RRB n=135, SC>RRB n=11). Abbreviations: FIQ = fullscale IQ; ADI-R = Autism Diagnostic Interview Revised; ADOS = Autism Diagnostic Observation Schedule; SC = social-communication; RRB = restricted repetitive behaviors; SA = social affect; CSS = calibrated severity score. ^ii^ DSM-5 domain percentage scores used for the SC-RRB difference score computation.

|  | Discovery |  |  |  | Replication |  |  |  |
| --- | --- | --- | --- | --- | --- | --- | --- | --- |
|  | SC>RRB | SC=RRB | RRB>SC | F (p-val) | SC>RRB | SC=RRB | RRB>SC | F (p-val) |
| **N**<br>**(male)** | 137<br>(109) | 611<br>(468) | 141<br>(115) | - | 124<br>(97) | 629<br>(482) | 137<br>(115) | - |
| **Age in**<br>**years** | 7.24<br>(5.50) | 9.03<br>(5.38) | 10.05<br>(4.02) | 10.61<br>(2.77e-5) | 6.92<br>(5.14) | 9.10<br>(5.54) | 9.76<br>(4.35) | 10.80<br>(2.31e-5) |
| **ADI-R**<br>**SC<sup>ii</sup>** | 0.51<br>(0.13) | 0.31<br>(0.14) | 0.21<br>(0.10) | 212.57<br>(2.2e-16) |  |  |  | 191.67<br>(2.2e-16) |
| **ADI-R**<br>**RRB<sup>ii</sup>** | 0.21<br>(0.10) | 0.32<br>(0.14) | 0.49<br>(0.12) | 174.19<br>(2.2e-16) |  |  |  | 184.74<br>(2.2e-16) |
| **ADOS**<br>**SA**<br>**CSS<sup>i</sup>** | 7.46<br>(1.60) | 6.92<br>(2.01) | 6.11<br>(2.49) | 2.85<br>(0.06) | 6.96<br>(1.73) | 6.93<br>(2.04) | 6.62<br>(2.39) | 1.83<br>(0.16) |
| **ADOS**<br>**RRB**<br>**CSS<sup>i</sup>** | 8.37<br>(1.31) | 7.68<br>(2.30) | 7.37<br>(2.29) | 0.09<br>(0.91) | 7.42<br>(2.55) | 7.40<br>(2.30) | 6.62<br>(1.06) | 0.43<br>(0.64) |
| **FIQ<sup>i</sup>** | 106.83<br>(17.10) | 103.89<br>(18.97) | 102.69<br>(14.59) | 0.32<br>(0.72) | 117.27<br>(13.76) | 105.44<br>(17.96) | 106.92<br>(15.88) | 2.38<br>(0.09) |

### Subtyping Analyses

To label subtypes by SC-RRB balance, we first computed difference scores between SC and RRB to estimate the level of SC-RRB balance, whereby values above 0 indicate an individual with higher SC versus RRB (SC>RRB), whereas values below 0 indicate the reverse (RRB>SC). These SC-RRB difference scores were then z-normalized using the mean and standard deviation estimated separately for Discovery and Replication sets. A z-score cutoff was used to derive subtype labels. Individuals falling above the z-cutoff (e.g., z>1) were labeled as SC>RRB, while individuals falling below the negative value of the z-cutoff (e.g., z<-1) were labeled as RRB>SC. All individuals between the z-cutoffs were considered SC=RRB. Because the choice of a z-cutoff is arbitrary, we ran all analyses across a range of z-thresholds from z=0.5 to z=1, in steps of 0.1. This approach allows us to report results across thresholds rather than using only one arbitrarily defined threshold. For the later functional connectivity analyses, this approach also allowed us to identify a consensus result which is consistent irrespective of the z-threshold used to label subtypes. To make out-of-sample predictions, we used the mean and SD norms estimated from the NDAR Discovery set to z-score and label individuals in the NDAR Replication set. These predicted labels on the Replication set were then compared to the actual subtype labels computed using the mean and SD norms derived from the Replication set itself. To make subtype predictions in the EU-AIMS LEAP dataset, we combined both NDAR Discovery and Replication datasets into one large dataset. From this dataset, norms for the mean and standard deviation were computed (mean = 0.01045243, SD = 0.19482749) and used for the z-scoring procedure. SC-RRB difference z-scores were then computed and a z-threshold was applied to the EU-AIMS LEAP dataset to generate subtype labels.

In addition to this SC-RRB difference z-score subtyping approach, we also used other unsupervised clustering methods for identifying subtypes. These methods utilize agglomerative hierarchical clustering with Euclidean distance and the ward.D2 method. The optimal number of clusters was determined by a majority vote of 23 different metrics for determining the optimal number of clusters (e.g., using the *NbClust* library in *R*)^27^. With another approach, we ran the same hierarchical clustering analyses, but cut dendrograms to define subtypes using a dynamic hybrid tree cut algorithm, as utilized in past work^28,29^.

### EU-AIMS LEAP Dataset

The EU-AIMS LEAP data comes from a large multisite European initiative with the aim of identifying biomarkers for ASD^19^. In this study, EU-AIMS LEAP data is utilized to examine how SC-RRB balance subtypes may differ in intrinsic functional connectomic organization using rsfMRI data. rsfMRI data from EU-AIMS LEAP has been analyzed for case-control differences in prior work^21,30^. EU-AIMS LEAP recruited 437 individuals with ASD and 300 TD individuals, both male and female, aged between 6 and 30 years. Participants underwent comprehensive clinical, cognitive, and MRI assessment at one of the following five centers: Institute of Psychiatry, Psychology and Neuroscience, King’s College London, United Kingdom; Autism Research Centre, University of Cambridge, United Kingdom; Radboud University Nijmegen Medical Centre, the Netherlands; University Medical Centre Utrecht, the Netherlands; and Central Institute of Mental Health, Mannheim, Germany. The study was approved by the local ethical committees of participating centers, and written informed consent was obtained from all participants or their legal guardians (for participants <18 years). For further details about the study design, we refer to Loth et al.,^20^, and for a comprehensive clinical characterization of the LEAP cohort, we refer to Charman et al.,^19^. In the present study, we selected all participants for whom structural and rsfMRI data were available. However, n=120 participants had to be excluded from the analysis because of missing ADI-R item-level data (n=64), missing IQ data (n=3), or because preprocessing could not be completed for a variety of reasons (e.g., registration/normalization errors because of poor quality MPRAGE data, poor anatomical coverage, or large anatomical deviance such as large ventricles (n=39), incomplete rsfMRI data (n=3), errors in convergence of the ME-ICA algorithm (n=11)). The final sample size was n=266 autistic and n=243 TD participants. This final sample was split into independent Discovery and Replication sets (balanced for sex and age) for the purpose of identifying functional connectivity differences that are replicable. As an example of sample sizes once split into autism subtypes at a z-threshold of 1, within the Discovery set there were n=77 SC=RRB, n=50 SC>RRB, and n=121 TD individuals. Within the Replication set there were n=83 SC=RRB, n=49 SC>RRB, and n=122 TD individuals. N=7 (n=6 Discovery, n=1 Replication) were classified as RRB>SC and because the sample sizes were too small, we did not analyze this subtype further for functional connectivity differences. We tested subtypes on a variety of different phenotypic measures including the ADOS-2, Social Responsiveness Scale (SRS-2), Repetitive Behavior Scale (RBS-R), Short Sensory Profile (SSP) and the Vineland Adaptive Behavior Scales (VABS). See Table 2 for participant characteristics.

**Table 2:** Participant characteristics from the EU-AIMS LEAP dataset. At a z-threshold of 1, this table shows sample sizes and descriptive statistics alongside F-statistic and p-value from an ANOVA testing for an effect of group. For ADI-R, ADOS, SRS, RBS, SSP, and VABS, the F-statistic and p-value refer to a group difference between SC=RRB vs SC>RRB, while for age, mean FD, and FIQ, the F-statistic and p-value refer to a model that takes into account all groups. ^i^ Sample sizes: ADOS (Discovery, RRB>SC n=6, SC=RRB n=76, SC>RRB n=48; Replication RRB>SC n=1, SC=RRB n=81, SC>RRB n=46); SRS (Discovery, RRB>SC n=3, SC=RRB n=67, SC>RRB n=44, TD n=68; Replication RRB>SC n=1, SC=RRB n=76, SC>RRB n=41, TD n=65); RBS (Discovery, RRB>SC n=3, SC=RRB n=64, SC>RRB n=43, TD n=68; Replication RRB>SC n=1, SC=RRB n=73, SC>RRB n=41, TD n=63); SSP (Discovery, RRB>SC n=2, SC=RRB n=44, SC>RRB n=33, TD n=59; Replication RRB>SC n=1, SC=RRB n=50, SC>RRB n=32, TD n=54); Vineland (Discovery, RRB>SC n=4, SC=RRB n=74, SC>RRB n=45, TD n=34; Replication RRB>SC n=1, SC=RRB n=76, SC>RRB n=44, TD n=39). Abbreviations: FD = framewise displacement; FIQ = full-scale IQ; ADI-R = Autism Diagnostic Interview Revised; ADOS = Autism Diagnostic Observation Schedule; SC = social-communication; RRB = restricted repetitive behaviors; SA = social affect; CSS = calibrated severity score; SRS = Social Responsiveness Scale; RBS = Repetitive Behavior Scale; SSP = Short Sensory Profile; VABS = Vineland Adaptive Behavior Scales; Comm = Communication; DLS = Daily Living Skills; Soc = Socialization; ABC = Adaptive Behavior Composite. ^ii^ DSM-5 domain percentage scores used for the SC-RRB difference score computation.

|  | Discovery |  |  |  |  | Replication |  |  |  |  |
| --- | --- | --- | --- | --- | --- | --- | --- | --- | --- | --- |
|  | SC>RRB | SC=RRB | RRB>SC | TD | F (p-val) | SC>RRB | SC=RRB | RRB>SC | TD | F (p-val) |
| **N (male)** | 50 (36) | 77 (59) | 6 (3) | 121 (80) | - | 49 (36) | 83 (60) | 1 (1) | 122 (75) | - |
| **Age in years** | 16.26 (5.03) | 16.29 (5.72) | 18.08 (7.13) | 16.83 (5.23) | 0.34 (0.79) | 16.30 (5.21) | 16.51 (5.67) | 11.45 (N/A) | 16.86 (6.07) | 0.63 (0.59) |
| **FIQ** | 98.27 (18.28) | 98.94 (18.89) | 102.67 (14.63) | 105.72 (18.33) | 2.34 (0.07) | 95.43 (20.11) | 103.28 (16.55) | 148.00 (N/A) | 104.94 (17.14) | 5.10 (0.001) |
| **Mean FD** | 0.23 (0.24) | 0.28 (0.45) | 0.29 (0.42) | 0.18 (0.15) | 2.17 (0.09) | 0.23 (0.21) | 0.25 (0.27) | 0.38 (N/A) | 0.23 (0.46) | 0.10 (0.95) |
| **ADI-R SC<sup>ii</sup>** | 0.46 (0.13) | 0.31 (0.14) | 0.18 (0.12) | - | 29.19 (3.70e-11) | 0.51 (0.13) | 0.28 (0.13) | 0.15 (N/A) | - | 54.60 (<0.001) |
| **ADI-R RRB<sup>ii</sup>** | 0.16 (0.09) | 0.29 (0.16) | 0.47 (0.09) | - | 24.87 (7.63e-10) | 0.20 (0.13) | 0.23 (0.13) | 0.40 (N/A) | - | 1.98 (0.14) |
| **ADI-R Soc** | 18.52 (6.44) | 16.83 (6.81) | 15.83 (8.33) | - | 2.70 (0.10) | 19.39 (5.87) | 14.96 (5.92) | 16.00 (N/A) | - | 19.47 (2.15e-5) |
| **ADI-R Comm** | 14.54 (5.03) | 14.00 (5.87) | 14.67 (6.31) | - | 0.94 (0.33) | 15.61 (4.64) | 11.84 (5.36) | 7.00 (N/A) | - | 19.29 (2.34e-5) |
| **ADI-R RRB** | 2.96 (2.02) | 5.08 (2.54) | 9.00 (1.26) | - | 26.72 (9.29e-7) | 3.88 (2.54) | 3.94 (2.29) | 6.00 (N/A) | - | 0.001 (0.97) |
| **ADOS SA CSS<sup>i</sup>** | 6.31 (2.54) | 6.22 (2.60) | 3.83 (3.37) | - | 0.06 (0.80) | 6.20 (2.98) | 5.68 (2.51) | 5.00 (N/A) | - | 1.48 (0.22) |
| **ADOS RRB CSS<sup>i</sup>** | 4.71 (2.79) | 4.86 (2.83) | 4.83 (4.22) | - | 0.20 (0.65) | 4.50 (2.83) | 4.91 (2.52) | 1.00 (N/A) | - | 1.61 (0.20) |
| **SRS<sup>i</sup>** | 73.23 (10.97) | 72.04 (12.35) | 71.33 (16.65) | 47.84 (9.40) | 0.01 (0.90) | 76.61 (10.61) | 67.13 (11.61) | 60.00 (N/A) | 47.23 (9.34) | 0.10 (0.75) |
| **RBS<sup>i</sup>** | 14.72 (13.27) | 18.83 (16.40) | 17.33 (10.69) | 2.15 (4.74) | 1.93 (0.16) | 21.59 (15.81) | 13.37 (11.30) | 13.00 (N/A) | 3.08 (11.54) | 12.37 (6.35e-4) |
| **SSP<sup>i</sup>** | 138.64 (28.33) | 135.86 (31.26) | 143.00 (33.94) | 177.86 (12.71) | 0.67 (0.41) | 134.62 (24.70) | 142.44 (26.14) | 153.00 (N/A) | 174.93 (19.38) | 2.33 (0.13) |
| **VABS Comm<sup>i</sup>** | 69.29 (16.26) | 73.65 (17.51) | 86.50 (20.68) | 91.97 (25.44) | 1.83 (0.17) | 69.50 (14.57) | 82.01 (13.81) | 99.00 (N/A) | 92.74 (25.45) | 21.97 (7.66e-6) |
| **VABS DLS<sup>i</sup>** | 68.31 (16.18) | 73.70 (16.61) | 71.00 (9.63) | 90.74 (20.28) | 3.31 (0.07) | 68.27 (15.83) | 78.24 (15.27) | 74.00 (N/A) | 91.10 (22.65) | 12.26 (6.60e-4) |
| **VABS Soc<sup>i</sup>** | 66.24 (15.96) | 70.78 (15.69) | 70.75 (10.34) | 96.21 (23.67) | 2.82 (0.09) | 63.43 (15.88) | 75.91 (15.22) | 76.00 (N/A) | 98.90 (27.04) | 18.79 (3.13e-5) |
| **VABS ABC<sup>i</sup>** | 65.71 (15.45) | 71.27 (13.20) | 74.00 (12.03) | 92.06 (23.04) | 4.44 (0.03) | 65.25 (13.82) | 77.09 (12.83) | 81.00 (N/A) | 93.00 (25.39) | 24.02 (3.16e-6) |

### EU-AIMS LEAP fMRI Data Acquisition

MRI data were acquired on 3T scanners: General Electric MR750 (GE Medical Systems, Milwaukee, WI, USA) at Institute of Psychiatry, Psychology and Neuroscience, King’s College London, United Kingdom (KCL); Siemens Magnetom Skyra (Siemens, Erlangen, Germany) at Radboud University Nijmegen Medical Centre, the Netherlands (RUMC); Siemens Magnetom Verio (Siemens, Erlangen, Germany) at the University of Cambridge, United Kingdom (UCAM); Philips 3T Achieva (PhilipsHealthcare Systems, Best, The Netherlands) at University Medical Centre Utrecht, the Netherlands (UMCU); and Siemens Magnetom Trio (Siemens, Erlangen, Germany) at Central Institute of Mental Health, Mannheim, Germany (CIMH). Procedures were undertaken to optimize the MRI sequences for the best scanner-specific options, and phantoms and travelling heads were employed to assure standardization and quality assurance of the multisite image-acquisition^19^. Structural images were obtained using a 5.5 minute MPRAGE sequence (TR=2300ms, TE=2.93ms, T1=900ms, voxels size=1.1×1.1×1.2mm, flip angle=9°, matrix size=256×256, FOV=270mm, 176 slices). An eight-to-ten minute resting-state fMRI (rsfMRI) scan was acquired using a multi-echo planar imaging (ME-EPI) sequence^31,32^; TR=2300ms, TE~12ms, 31ms, and 48ms (slight variations are present across centers), flip angle=80°, matrix size=64×64, in-plane resolution=3.8mm, FOV=240mm, 33 axial slices, slice thickness/gap=3.8mm/0.4mm, volumes=200 (UMCU), 215 (KCL, CIMH), or 265 (RUMC, UCAM). Participants were instructed to relax, with eyes open and fixate on a cross presented on the screen for the duration of the rsfMRI scan.

### EU-AIMS LEAP fMRI Preprocessing

Multi-echo rsfMRI data were preprocessed with the multi-echo independent components analysis (ME-ICA) pipeline, implemented with the *meica* python library (v3.2) (https://github.com/ME-ICA/me-ica). ME-ICA implements both basic fMRI image preprocessing and decomposition-based denoising that is specifically tailored for multi-echo EPI data. For the processing of each subject, first the anatomical image was skull-stripped and then warped nonlinearly to the MNI anatomical template using AFNI 3dQWarp. The warp field was saved for later application to functional data. For each functional dataset, the first TE dataset was used to compute parameters of motion correction and anatomical-functional coregistration, and the first volume after equilibration was used as the base EPI image. Matrices for de-obliquing and six-parameter rigid body motion correction were computed. Then, 12-parameter affine anatomical-functional coregistration was computed using the local Pearson correlation (LPC) cost functional, using the gray matter segment of the EPI base image computed with AFNI 3dSeg as the LPC weight mask. Matrices for de-obliquing, motion correction, and anatomical-functional coregistration were combined with the standard space nonlinear warp field to create a single warp for functional data. The dataset of each TE was then slice-time corrected and spatially aligned through application of the alignment matrix, and the total nonlinear warp was applied to the dataset of each TE. No time-series filtering was applied in the preprocessing phase. No spatial smoothing was applied during preprocessing.

The preprocessed multi-echo time-series datasets were then used by the ME-ICA pipeline to leverage information in the multiple echoes to compute an optimal weighting of TE at each voxel^33^, producing an ‘optimally combined’ time-series dataset. This optimal combination procedure has been shown to double temporal signal-to-noise ratio (tSNR) over traditional single echo EPI data^34^. This preprocessed optimally combined time-series dataset was then fed into a denoising procedure based on independent components analysis (ICA) and scoring components by ρ and κ pseudo-F statistics that indicate degree of TE-independence or TE-dependence. Components with high ρ and low κ are components high in non-BOLD related contrast (i.e. nonBOLD artefact signal), while components with high κ and low ρ indicate components high in BOLD-related contrast. ME-ICA identifies in an automated fashion high ρ and low κ non-BOLD components and removes them from the optimally combined time-series dataset to produce the final multi-echo denoised dataset. This procedure has been shown to be very effective in removing various types of non-BOLD artefact from rsfMRI data, including head motion artefact, flattens DVARS traces induced by head motion, and increases tSNR by a factor of 4 over and above traditional single echo EPI data^31,32,34,35^. The final multi-echo denoised datasets were used in further connectivity analyses. Head motion estimates and DVARS were estimated in order to show the impact of denoising on reducing non-BOLD artefact due to head motion (see Fig. S1 for examples). In the EU-AIMS LEAP data, groups did not differ in mean FD (see Table 2).

### EU-AIMS LEAP Functional Connectivity Analyses

To assess large-scale intrinsic functional organization of the brain we input the multi-echo denoised data into a group-ICA analysis. Dual regression was then utilized to back-project spatial maps and individual time-series for each component and subject. Both group-ICA and dual regression were implemented with FSL’s MELODIC and Dual Regression tools (www.fmrib.ox.ac.uk/fsl). For group-ICA, we constrained the dimensionality estimate to 30. Of the 30 final components, 11 were discarded after visual examination of spatial maps which indicated that they did not correspond to well-known rsfMRI networks and instead resembled white matter or other artefacts^36^. See Fig. S2 for visual depiction of the 19 ICs used in further analysis.

Time courses for each subject and each independent component (IC) were used to model between-component connectivity. This was achieved by constructing a partial correlation matrix amongst all 19 components using Tikhonov-regularization (i.e. ridge regression, rho=1) as implemented within the *nets_netmats.m* function in the FSLNets MATLAB toolbox (https://fsl.fmrib.ox.ac.uk/fsl/fslwiki/FSLNets). The aim of utilizing partial correlations was to estimate direct connection strengths in a more accurate manner than can be achieved with full correlations, which allow more for indirect connections to influence connectivity strength^37–39^. Partial correlations were then converted into Z-statistics using Fisher’s transformation for further statistical analyses. The lower diagonal of each subject’s partial correlation matrix was extracted for a total of 171 separate component-pair comparisons.

To identify replicable subtype effects on functional connectivity, we partitioned the EU-AIMS LEAP dataset into Discovery and Replication sets. This was achieved via a random half split of the subtypes within each scanning site and balancing for sex. TD comparison groups for Discovery and Replication sets were also selected via a random split balancing sex and achieving an age-match (achieved using the *MatchIt* library in R with the default method of nearest neighbor matching). Models implementing the main hypothesis tests of subtype differences were computed as linear mixed effect models (*lme* function from the *nlme* library in R), whereby connectivity was the dependent variable, and subtype, sex, and age were used as fixed effect independent variables and site was modeled with random intercepts as a random effect. These models were computed separately for the Discovery and Replication set. Connectivity pairs were deemed as showing replicable subtype differences if the Discovery set showed an effect at p<0.05 and the replication Bayes Factor statistic^40^ computed on t-statistics from Discovery and Replication sets was greater than 10 *(repBF>10*), indicating strong evidence in favor of replication.

Because subtyping depends on the choice of a z-threshold, we ran the connectivity analyses across a range of z-thresholds from z=0.5 to z=1, moving up in steps of 0.1. Across all these z-thresholds, we identified ‘consensus edges’, defined as replicable subtype connectivity differences that appear at all z-thresholds. These edges are focused since they are the robust subtype connectivity differences that are not dependent on a particular z-threshold for labeling the subtypes. For each threshold, we also counted up the number of edges that are common across subtypes and with similar directionality in order to estimate how often subtypes show similar functional connectivity differences.

To contrast the subtyping approach to a more dimensional approach where z-normalized SC-RRB differences scores are left continuous, we also ran similar mixed effect models where these continuous scores are the primary independent variable of interest rather than the subtype variable. Because the z-normalized difference score does not capture overall severity level well (e.g., an individual with low SC and RBB has a difference near 0 just like an individual with high SC and RRB), we also ran models whereby continuous SC or RRB scores were used as independent variables rather than the z-normalized difference score. This allowed for another contrast to test if overall level of severity within each domain could explain connectivity strength. In each of these dimensional models, the same criteria for identifying replicable effects in the subtype models was used (e.g., p<0.05 in the Discovery set and a *repBF* > 10).

### Gene expression decoding analyses

To identify genes whose spatial expression pattern is similar to subtype-specific ICs, we used the Gene Expression Decoding feature embedded within Neurosynth^23^ to identify genes that are statistically similar in their expression profile in a consistent manner across all 6 donor brains within the Allen Institute Human Brain Atlas^22^. The analysis first utilizes a linear model to compute similarity between the observed rsfMRI IC map and spatial patterns of gene expression for each of the six brains in the Allen Institute dataset. The slopes of these subject-specific linear models encode how similar each gene’s spatial expression pattern is with our rsfMRI IC maps. These slopes were then subjected to a one-sample t-test to identify genes whose spatial expression patterns are consistently of high similarity across the donor brains to the rsfMRI IC maps we input. This analysis was restricted to cortical tissue since all of the networks being analyzed are primarily cortical. The resulting list of genes was then thresholded for multiple comparisons and only the genes surviving FDR q < 0.05 and also had a positive t-statistic value were considered.

### Enrichment analyses with autism-associated gene lists

To test if network-associated genes were enriched for different classes of autism-associated genes we first curated a list of genes known at genetic and transcriptomic levels to be associated with autism. At the genetic level, we utilized the list of 102 genes reported by Satterstrom et al.,^41^ that are rare de novo protein truncating variants that are associated with a diagnosis of autism (ASD dnPTV). A second list of autism-associated genes (ASD SFARI) at the genetic level was the list curated by SFARI Gene (https://gene.sfari.org/). We utilized the entire list of genes in categories S, 1, 2, and 3 for these analyses (downloaded on July 16, 2020). At the transcriptomic level we used several lists. First, we used a list of differentially expressed genes in autism postmortem frontal and temporal cortex tissue from Gandal et al.,^42^ and this list was further split by genes that were downregulated (ASD DE Downreg) or upregulated (ASD DE Upreg) in autism. To contrast these enrichments with other psychiatric diagnoses that are genetically correlated with autism, we also use differentially expressed genes in schizophrenia (SCZ DE) and bipolar disorder (BD DE), identified from the same Gandal et al., study^42^. To go beyond differentially expressed genes in bulk tissue samples, we also examined autism differentially expressed genes identified in specific cell types - particularly, excitatory (ASD Excitatory) and inhibitory (ASD Inhibitory) neurons, microglia (ASD Microglia), astrocytes (ASD Astrocyte), oligodendrocytes (Oligodendrocyte), and endothelial (ASD Endothelial) cells^43^. Beyond differentially expressed genes, we utilized all genes identified in frontal and temporal cortical tissue that were members of co-expression modules identified to be downregulated (ASD CTX Downreg CoExpMods) or upregulated (ASD CTX Upreg CoExpMods) in autism^44^. All tests of enrichment were conducted with custom code written in R that computes enrichment odds ratios and p-values based on hypergeometric tests. The background total for these enrichment tests was set to 20,787, which is the total number of genes considered by the gene expression decoding analysis in Neurosynth. FDR was computed amongst all of the enrichment tests done and only tests that survived FDR q < 0.05 were interpreted further as statistically significant enrichments.

### Data and code availability

Tidy data and reproducible analysis code for this study is available at https://github.com/IIT-LAND/adir_subtyping.

## Results

### Highly accurate out-of-sample prediction of SC-RRB balance subtypes

In our first set of analyses, we sought to develop a model to predict ADI-R SC-RRB balance subtypes from the NDAR datasets. Relatively equal Discovery (n=889) and Replication (n=890) datasets were partitioned from the total n=1,781 individuals in NDAR and this split into Discovery and Replication was balanced as a function of the originating datasets and sex. Using z-normalized difference scores, we split the dataset into SC=RRB, SC>RRB, and RRB>SC subtypes (Fig. 2). Importantly, the subtype labels were first defined separately in Discovery and Replication sets based on the statistical norms (i.e. mean and SD) estimated on each set. This ensures that the definition of the labels in each set is done independently of the other. If the statistical norms for the computation of z-normalized difference scores (e.g., mean and SD) are highly similar between Discovery and Replication, then the subtyping model learned from the Discovery set will likely be highly generalizable and produce high accuracy values in the Replication set. However, if the statistical norms are highly different between Discovery and Replication, the model learned from Discovery data will not generalize well to the labels in the Replication set and would thus produce poor out-of-sample prediction accuracy.

**Fig. 2:**
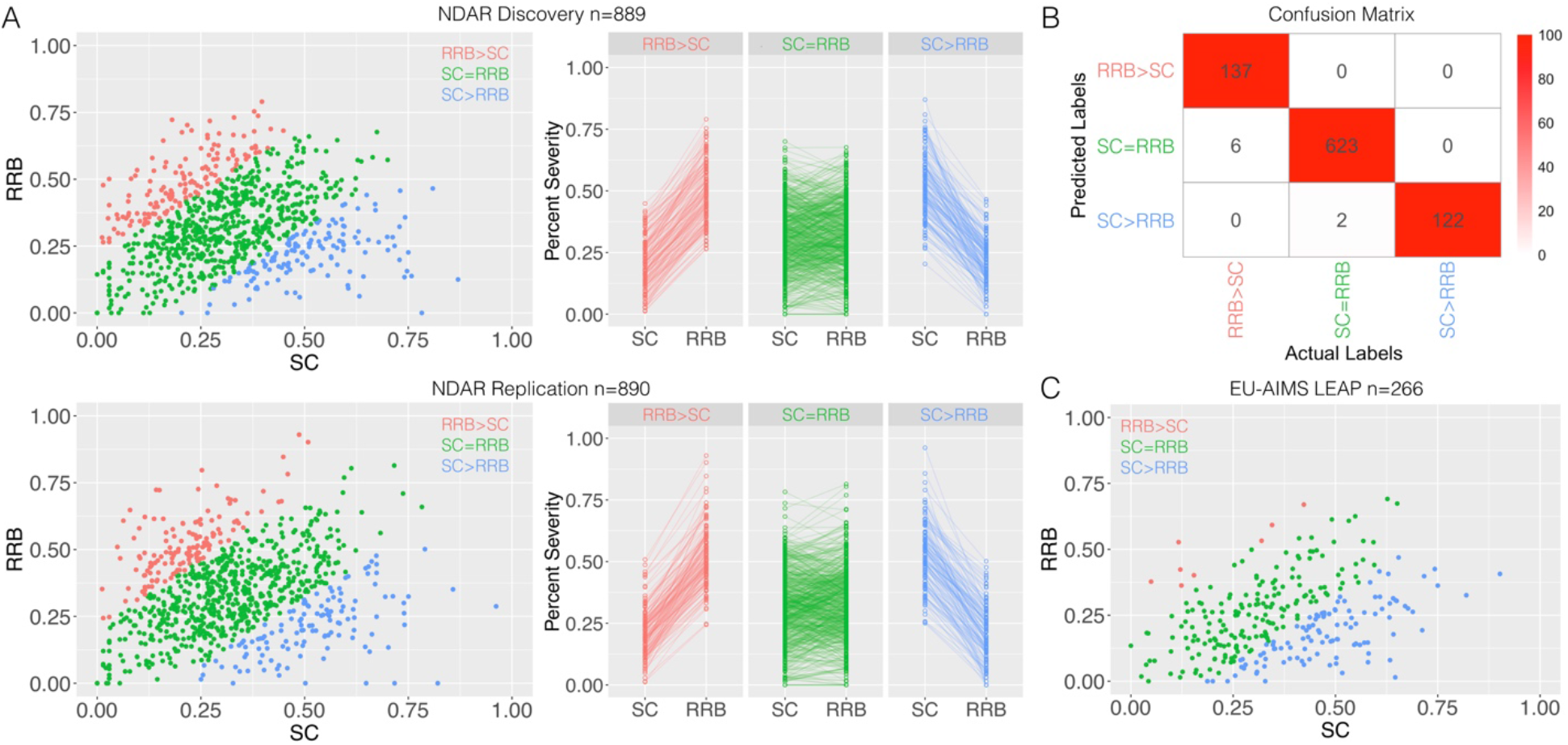
Supervised subtyping of autism by SC-RRB balance. Panel A shows the subtypes derived from a z-normalized difference score of SC-RRB, with a z-score threshold for cutting the subtypes at z = 1. Red shows the RRB>SC subtype, green shows the SC=RRB subtype, and blue shows the SC>RRB subtype. Panel B shows a confusion matrix with actual subtype labels for the NDAR Replication dataset along the columns and the subtyping model’s predicted labels (trained on the NDAR Discovery dataset) along the rows. The colors within the cells indicate the percentage of individuals relative to the actual labels with predicted labels in each cell. Over a range of z-thresholds from 0.5 to 1, the accuracy ranged from 97-99% accuracy. Panel C shows SC-RRB subtypes from the EU-AIMS LEAP datasets derived using norms estimated from NDAR.

Applying this approach across a range of z-thresholds (e.g., z = 0.5 up to z =1 in steps of 0.1), we find that out-of-sample prediction accuracy is very high (e.g., 97-99%). The reason for this high accuracy is visually evident in the high degree of similarity in distributions of Discovery and Replication sets in the scatterplots in Fig. 2A. Examination of sex across these subtypes did not yield any significant between-subtype differences (Discovery: *χ*^2^ = 1.91; *p* = 0.38; Replication: *χ*^2^ = 3.50; *p* = 0.17), with a 3:1 to 5:1 sex ratio of males to females. Subtypes did differ in age at the time of ADI-R interview, with the SC>RRB group being younger than the other subtypes (Discovery: *F(2,886)* = 10.61, *p* = 2.77e-5; Replication: *F(2,887)* = 10.80, *p* = 2.31e-5). See Table 1 for descriptive statistics.

Contrasting this z-score approach to subtyping with unsupervised clustering methods (Fig. S3) that use static tree cut heights along with internal cluster validation metrics for choosing the optimal number of clusters, we found that such SC-RRB balance subtypes are not easily identifiable in a consistent fashion across Discovery and Replication cohorts with such blind methods. However, when using an automated dynamic hybrid tree-cutting algorithm that adaptively modifies cutting the dendrogram at different heights^28,29^, we are able to get relatively close to finding similar partitions in Discovery (6 subtypes) versus Replication (7 subtypes) sets (Fig. S4).

### Replicable subtype-specific functional connectivity differences

We next evaluated whether such SC-RRB balance subtypes are differentiated from TD comparison groups in rsfMRI functional connectivity. Because subtypes are defined based on thresholding the z-normalized SC-RRB difference score, we identified ‘consensus edges’ as functional connectivity differences between the autism subtype versus TD that consistently appear across every z-threshold examined. Fig. 3 summarizes the consensus edges in each subtype for both the LEAP Discovery and Replication sets. Relative to the TD group, the SC=RRB subtype is characterized by on-average hyperconnectivity between the anterior salience network (IC07) and a medial motor network (IC13) (effect sizes at z=1 threshold: Discovery *Cohen’s d* = 0.36; Replication *Cohen’s d* = 0.51; *repBF* = 390) as well as hypoconnectivity between visual association (IC03) and somatomotor (IC12) networks (effect sizes at z=1 threshold: Discovery *Cohen’s d* = −0.41; Replication *Cohen’s d* = −0.36; *repBF* = 16). The somatomotor network was also hypoconnected in SC>RRB relative to TD, but with the bilateral perisylvian (IC17) network (effect sizes at z=1 threshold: Discovery *Cohen’s d* = −0.40; Replication *Cohen’s d* = −0.41; *repBF* = 23). In contrast to comparing autism subtypes to TD, we also directly compared SC=RRB versus SC>RRB. This analysis did not yield any significant replicable differences, indicating that while these subtypes can replicably differ relative to a TD comparison group in qualitatively unique ways, the difference between each other may not be replicably large enough to detect at current sample sizes (effect sizes for z=1 threshold: IC07-IC13, Discovery *Cohen’s d* = 0.10, Replication *Cohen’s d* = 0.01; IC03-IC12, Discovery *Cohen’s d* = −0.10, Replication *Cohen’s d* = −0.13; IC12-IC17, Discovery *Cohen’s d* = 0.27, Replication *Cohen’s d* = 0.11). For the full set of statistical results at z=1 threshold across all comparisons see Supplementary Table 2. Thus, the connectivity results suggest a mixture of some overlap in affected networks in both subtypes (e.g., IC12), alongside some qualitative specificity of networks affected in only one of the subtypes (e.g., IC03, IC07, IC13 for SC=RRB; IC17 for SC>RRB). Importantly, this subtype-distinctiveness is subtle and relative to TD, but does not heavily differ when subtypes are directly compared to each other.

**Fig. 3:**
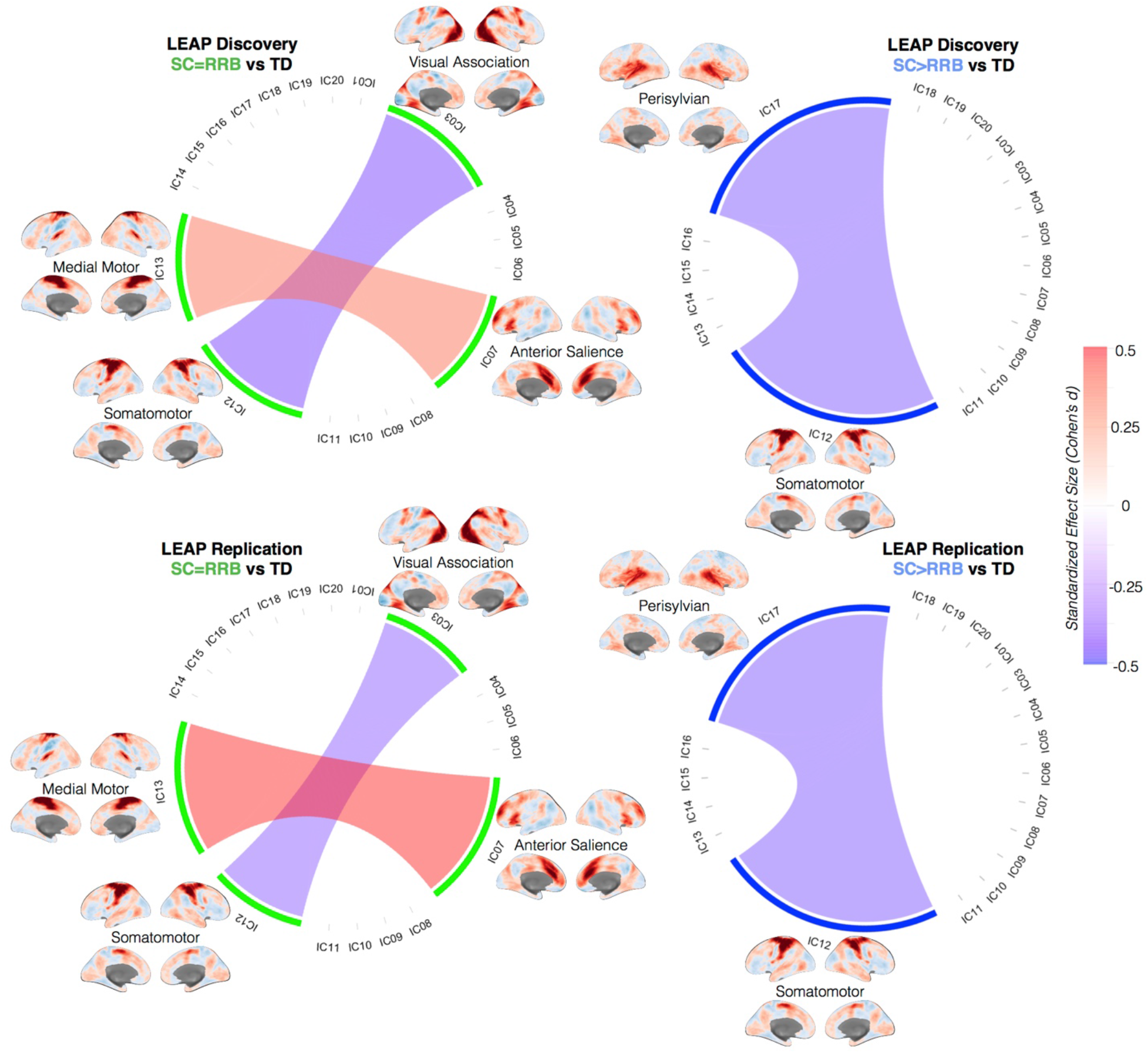
Replicable subtype differences in functional connectivity. This figure shows chord diagrams of replicable functional connectivity differences between SC=RRB vs TD (left) or SC>RRB vs TD (right), when subtypes are defined at a z-threshold of 1. However, edges shown in these diagrams are consensus edges that appear in every analysis of connectivity differences irrespective of the z-threshold used to define the subtypes. Red edges indicate hyperconnectivity in the autism subtype relative to TD, while blue edges indicate hypoconnectivity in autism subtype relative to TD. Intensity of edge color indicates standardized effect size (Cohen’s d) shown on the colorbar on the right. The cortical surface renderings of each component are unthresholded z-stat maps. Areas with higher z-stats (dark red) are of primary importance for the IC map. The top row shows effects in the EU-AIMS LEAP Discovery set, while bottom row shows effects in the EU-AIMS LEAP Replication set.

Because the subtyping approach uses the difference score between SC and RRB, this metric does not distinguish individuals by overall level of severity. For example, an individual with low SC and RRB severity is treated similarly to an individual with high SC and RRB severity. This leaves open the possibility that degree of severity on a continuum from high to low could also explain variability in functional connectivity. To test this hypothesis, we constructed a dimensional model to predict connectivity strength from SC or RRB severity as a continuous variable. However, there were no instances whereby SC or RRB severity as a continuous measure could replicably predict connectivity strength. Similarly, when using the z-normalized SC-RRB difference score as a continuous variable, we also found no replicable significant effects on connectivity. For the full set of statistics see Supplementary Table 2. These results provide a dimensional model contrast to the categorical subtyping approach and suggests that modeling continuous SC or RRB variability may be less sensitive as a predictor of functional connectivity compared to SC-RRB balance subtypes.

### Divergent functional genomic underpinnings of subtype-specific neural circuitry

In the next analysis, we asked if known autism-associated genes are enriched amongst genes known to be highly expressed in these subtype-associated rsfMRI networks. We first identified lists of genes whose spatial expression topology in the Allen Institute Human Brain Atlas^22^ is similar to rsfMRI connectivity networks^23^ that show replicable subtype differences. Once a set of genes are predicted to underpin such rsfMRI networks, we then asked whether those genes are highly overlapping with known sets of functional genomic mechanisms linked to autism (see Fig. 4A for a visual representation of the analysis approach and Supplementary Table 3 for the full set of gene lists used in these analyses). Underscoring functional genomic overlap between the subtypes, all networks except for IC07 were enriched for a variety of similar autism-associated gene lists – such as, high penetrance rare de novo protein truncating variants (IC03, IC17), genes associated with autism from the SFARI Gene database (IC03, IC12, IC17), genes and coexpression modules that are downregulated in expression (IC1, IC03, IC12, IC17), and genes differentially expressed in excitatory and inhibitory neurons (IC03, IC12, IC17) and astrocytes (IC17). Despite this overlap, our next analysis focused on genes that were specific to networks affecting only one of the subtypes. To achieve this aim, we removed genes that showed high levels of expression across multiple networks. The resulting lists are genes that are expressed specifically in only one of the networks affecting the SC-RRB imbalance subtypes. This analysis revealed that genes expressed SC>RRB-affected networks (i.e. specifically IC17) are enriched for SFARI ASD genes, autism-downregulated co-expression modules, and genes differentially expressed in excitatory and inhibitory neurons and astrocytes. In contrast, genes expressed within SC=RRB-affected networks (i.e. specifically IC03, IC13, and IC07) are enriched only for genes downregulated in expression in autism. Thus, much like the connectivity results, these results implicate a mixture of overlap as well as some specificity in the genomic mechanisms that can affect networks implicated in the subtypes.

**Fig. 4:**
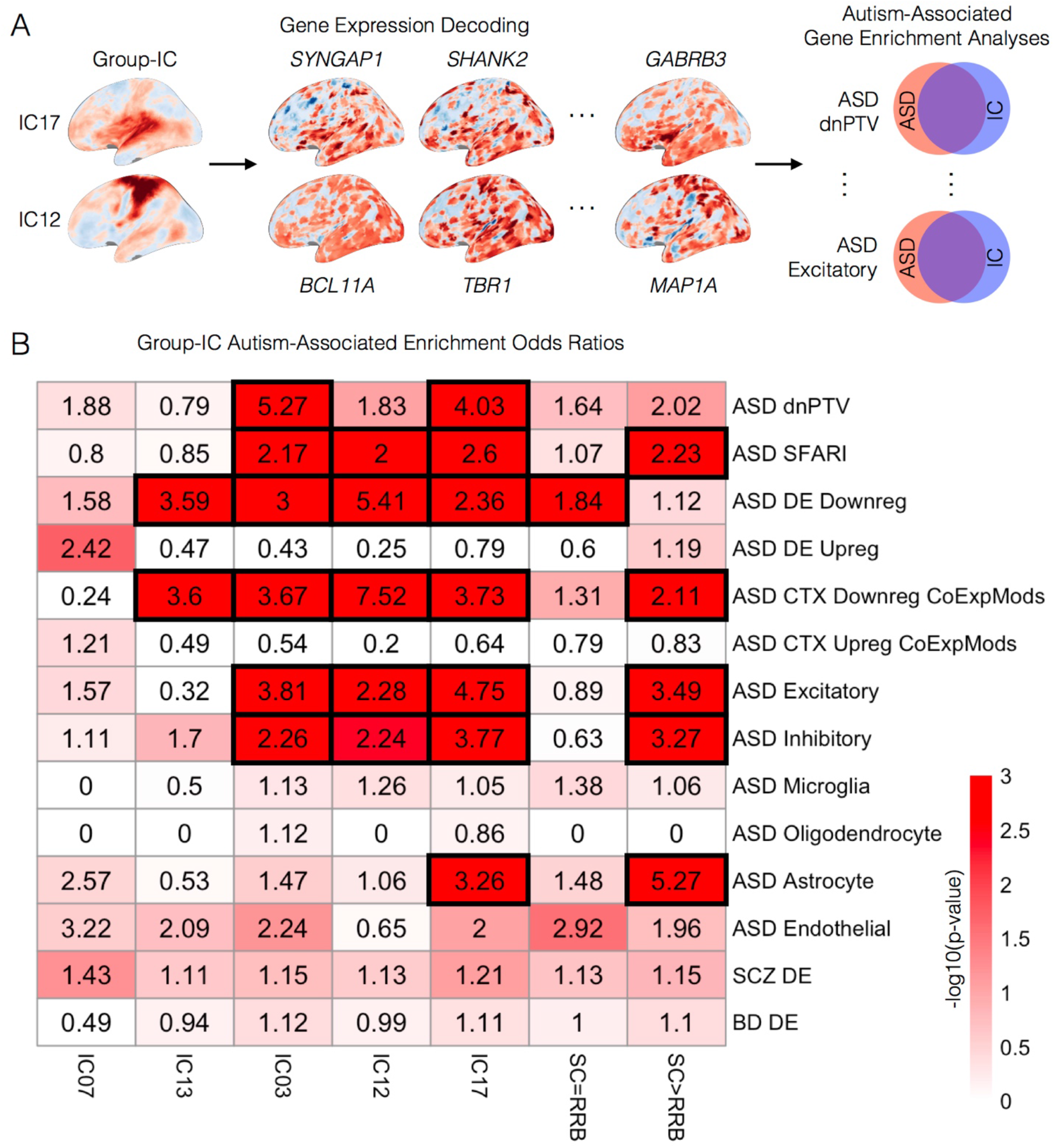
Overlap between genes expressed in functional connectivity networks and genes linked to autism. In panel A we depict the analysis approach of identifying genes which are highly expressed in similar spatial patterns to the rsfMRI spatial IC maps (i.e. gene expression decoding). Once IC gene lists have been identified, we test these lists for enrichment with known lists of autism-associated functional genomic mechanisms (top left). In panel B we show enrichment odds ratios (numbers in each cell) along with the −log10 p-value (coloring of the cells) for enrichment tests of specific networks (columns) against known lists of autism-associated genomic mechanisms (rows). Cells outlined with thick black rectangles survive FDR q<0.05. The column labeled SC>RRB shows the enrichment results when the gene list under consideration comprises genes unique to IC17, but not any of the other ICs. The column labeled SC=RRB shows the enrichment results when the gene list under consideration consists of genes unique to IC03, IC07, and IC13, but not IC12 or IC17. ASD dnPTV, Autism de novo protein truncating variants; ASD SFARI, SFARI Gene autism associated genes; ASD DE Downreg, autism differentially expressed downregulated genes; ASD DE Upreg, autism differentially expressed upregulated genes; ASD CTX Downreg CoExpMods, autism downregulated cortical co-expression modules; ASD CTX Upreg CoExpMods, autism upregulated cortical co-expression modules; ASD Excitatory, autism differentially expressed genes in excitatory neurons; ASD Inhibitory, autism differentially expressed genes in inhibitory neurons; ASD Microglia, autism differentially expressed genes in microglia; ASD Oligodendrocyte, autism differentially expressed genes in oligodendrocytes; ASD Astrocyte, autism differentially expressed genes in astrocytes; ASD Endothelial, autism differentially expressed genes in endothelial cells; SCZ DE, schizophrenia differentially expressed genes; BD DE, bipolar disorder differentially expressed genes.

## Discussion

In this work, we examined how autism SC-RRB balance subtypes are atypical at the level of macroscale neural circuitry measured with rsfMRI. Prior work has theorized that the core dyad of SC and RRB is fractionable at behavior, neural, and genetic levels^4–14^. However, it is unclear whether the road from genome to phenome (e.g., Fig. 1) is one that converges on a common pathway or is one of multiple pathways^15^. Here we find evidence consistent with a mixture of both common and multiple pathways. Consistent with the common pathway hypothesis, we first find no evidence of strong between-subtype differences in autism when subtypes are directly compared to each other. Rather, all replicable differences in functional connectivity appear when subtypes are compared to a TD comparison group. Also consistent with the idea of commonalities between the subtypes is the shared effect of somatomotor network hypoconnectivity with other networks. Gene expression decoding analysis of individual networks also showed some commonalities in enrichments with autism-associated gene lists.

Despite these commonalities, there was also evidence for subtle distinctions between the subtypes. First, the edges identified as replicably different between the subtypes relative to TD were never the same for SC=RRB and SC>RRB subtypes. This effect indicates that relative to the TD norm, subtle but replicable on-average differences in network connectivity exist. This highlights how even though functional neural circuitry organization appears to be mostly shared between the subtypes, each subtype diverges slightly but also uniquely from typical development in their functional organization. It is possible that these on-average subtype differences relative to TD are driven by a smaller subset of individuals within the subtypes with the most dramatic differences from TD. However, it is also possible that phenotypic variability within our subtypes might result not simply from distinct neurocircuitry abnormalities but from more complex and subtle combinations of shared and divergent neurocircuits affecting the balance between symptom domains. Second, upon isolating the genes expressed specifically within subtype-specific networks, we find a different pattern of autism-associated gene enrichment. Thus, rather than implicating a commonality in the genomic mechanisms that underpin different subtypes, this effect is consistent with the idea that some of the subtype-relevant genomic mechanisms differentially affect specific neural circuits such as perisylvian, visual association, or medial motor networks.

Regarding the importance of specific networks identified in our analyses, the somatomotor network (IC12) has been implicated in past work on autism. Somatosensory areas have been shown to be some of the most informative regions in prior case-control classifier studies using rsfMRI data^45^. Additionally, prior case-control analyses of the EU-AIMS LEAP dataset find that somatomotor areas show reduced degree centrality and autism-related hyperconnectivity with cerebellar networks^21,30^. However, the lack of somatomotor hypoconnectivity with visual association or perisylvian networks in prior case-control studies highlights the potential importance and added value of subtyping for revealing more subtle effects that can be masked with case-control contrasts. The perisylvian network that is hypoconnected with somatomotor circuitry in SC>RRB overlaps with a variety of areas implicated in early development of autism, particularly for auditory processing and language^46–50^. Integration of information processing between this network and others that play a role in embodied somatosensory and social cognitive processing^51,52^, such as the somatomotor network (IC12), could be important for explaining the more pronounced difficulties these individuals have within SC compared to RRB. Somatomotor hypoconnectivity with perisylvian auditory and visual association circuitry could also be important for pointing towards atypical multisensory integration that has been documented in autism^53,54^, particularly with regards to auditory-somatosensory^55^ and visual-somatomotor integration^56–60^.

In addition to somatomotor hypoconnectivity, we also observed a medial motor network which was hyperconnected to the anterior salience network in the SC=RRB subtype. The additional implication of another motor-relevant networks in this subtype underscores the importance of motor circuitry^61–63^ and visual-motor integration atypicalities in autism^56–60^. The anterior salience network has also been identified in prior case-control studies. In younger cohorts, anterior salience areas are hyperconnected^64,65^, while in older cohorts, hypoconnectivity is observed^66^. While age could be a factor in explaining the discrepant findings from prior work, it likely cannot explain the SC=RRB hyperconnectivity finding. Here we age-matched the groups and additionally included age as a covariate in the statistical model. EU-AIMS LEAP also samples from a wide age range from 6 to 30 years of age, enabling the sample to include younger and older ages covered by prior work. Thus, age may not be the only explanation for salience network hyperconnectivity. Rather, this work suggests that SC-RRB heterogeneity and the presence of this balanced subtype could also drive such effects in case-control comparisons, particularly if the sample is enriched with this particular subtype.

We also identified autism-relevant genomic underpinnings behind these subtype-specific rsfMRI networks. Genes specific to SC>RRB networks are enriched for a number of genomic mechanisms linked to autism such as genes differentially expressed in excitatory and inhibitory neurons and astrocytes, downregulated co-expression modules, and high-risk genetic mutations associated with autism. These genomic underpinnings suggest that specific neuronal cell types involved in cortical excitation-inhibition balance^67,68^ may be especially important for the SC>RRB subtype. This effect also partially corroborates evidence suggesting that excitatory neurons are affected in specific types of autistic individuals that differ in patterns of clinical severity^43^. In contrast, SC=RRB networks lacked similar kinds of enrichments, suggesting that differing functional genomic mechanisms may be linked to this subtype.

Another important finding from the current work is the absence of replicable connectivity effects in simplistic models that treat SC and RRB separately as continuous predictors. These findings suggest that continuous variation (i.e. severity) within each domain separately may not have large impact on explaining variation in functional connectivity. Rather, the SC-RRB balance subtyping approach of jointly considering the unique mixture of both SC and RRB within an individual as a means to then categorically split the autism population into subtypes, could be a more fruitful first pass approach for explaining connectivity differences. For example, other work also suggests that categorical factors may be mixed within participants in a mosaic fashion, whereby different individuals will have different mixtures of continuous differences along the factors^69^. This idea of a blend between both categorical and dimensional explanations for connectivity can be seen in work showing that etiologically distinct mechanisms known to cause autism result in continuous differentiation along a manifold landscape of functional connectivity^70^. Thus, further work might expand on categorical distinctions put forth by SC-RRB balance models to explain continuous variability within such subtypes.

There are certain limitations and caveats that need to be discussed. First, the threshold for the z-score cutoff to define subtypes could be viewed as arbitrary. However, to guard against this issue, we re-ran the analysis across a range of thresholds from z=0.5 to z=1 and showed effects that are robust to the threshold used to label the subtypes. Accuracy for out-of-sample predictions is also high regardless of the threshold. This effect occurs largely because the data distributions and statistics used for the z-normalization are highly similar across large NDAR Discovery and Replication sets. In a situation where the data distributions were not similar, this high out-of-sample prediction accuracy would not have been obtained or may have fluctuated more substantially at different thresholds. Thus, while the choice of a threshold may not be well defined, any choice within the range we have analyzed of z=0.5 to z=1, will yield highly consistent results that are not biased by the choice of a threshold. The fact that the data distributions and sample statistics used for the z-normalization were so similar across well-matched NDAR Discovery and Replication sets allows for high-confidence that the large NDAR dataset is likely very close in accurately estimating the true population parameters, and this allows for the high degree of replicability and robustness of the subtyping approach. Second, the distinctions between these subtypes are not demarcated by large categorical separations. As such, when we applied other canonical unsupervised clustering methods to the data, such methods were not able to consistently identify the same subtypes in independent datasets (Fig. S2). An automated dynamic hybrid treecutting method to apply to hierarchical clustering^28,29^ was however, close to deriving relatively similar subtypes across Discovery and Replication sets (Fig. S3). Future work could explore the utility of this approach and the consensus subtypes derived from independent datasets with this methodology. However, the lack of very large separations between the boundaries for different subtypes of autistic individuals likely means that a more nuanced and theory-driven approach may be more fruitful than blind unsupervised approaches. Third, the RRB>SC group was not highly prominent in the EU-AIMS LEAP cohort. This observation is likely due to the fact that NDAR includes studies that more heavily sample individuals from the population with higher RRB severity relative to EU-AIMS LEAP. For example, ADOS CSS scores for RRB are higher in NDAR than in LEAP (see Tables 1-2). Because NDAR pools from a much wider range of studies in different contexts compared to EU-AIMS LEAP, this could be an explanation for this difference. Fourth, direct comparisons of functional connectivity between SC=RRB and SC>RRB subtypes did not yield as large or replicable differences as when the subtypes are compared to TD. Thus, while there are unique consensus edges that appear when the autism subtypes are compared to TD, this result should not be taken to imply that the subtypes themselves are also highly different from each other. A likely reason for why these differences manifest when compared to TD but not when subtypes are compared directly may be due to effects driven by further subsets of individuals nested within the larger SC=RRB and SC>RRB subtypes. Alternatively, it could be that the SC-RRB subtyping approach does not allow for parsing apart the mechanisms that clearly distinguish different autistic individuals from each other. If autistic individuals are mosaics of many complex etiological mechanisms and those mechanisms have different effects on functional connectivity, then it may be that models quantifying such etiological mixtures may be better models of functional connectivity variation^69,70^. These individuals at the extremes of the functional connectivity distributions likely drive the on-average differences from TD. Future work that digs further into more granular divisions of the population may likely identify much larger differences when autism subtypes are compared directly. Fifth, we also discovered that dimensional models using continuous SC and RRB severity did not uncover any replicable associations with functional connectivity strength. This result could suggest that dimensional models that use continuous severity from the ADI-R are less effective than the subtyping approach. However, it could also be that dimensional models might be more sensitive with other measures of symptomatology (e.g., ADOS, SRS). Sixth, the subtyping here is based on the ADI-R. ADI-R is a commonly used ‘gold standard’ diagnostic instrument to aid clinical judgment regarding an autism diagnosis. However, other measures such as the ADOS could also have been used. For our purposes in this study, we chose to utilize the ADI-R over the ADOS due to the fact that participants come from a wide age range, and the ADOS would assess current symptomatology of the individual. If age has an effect on symptomatology^71–73^, then this could potentially bias the subtyping approach depending on the composition of the sample. On the other hand, because the ADI-R ‘diagnostic algorithm’ utilizes items that focus on early developmental and ‘most severe in lifetime’ symptomatology, we do not know how the individual might have changed across the lifespan of development. Additionally, it may be that measures of current symptomatology have a stronger association with measures of current functional connectivity than early childhood and lifetime snapshots of severity provided by the ADI-R. Future work that looks at how these ADI-R-derived SC-RRB balance subtypes might change over time would be informative from a developmental angle. It would also be important to investigate how observational measures such as the ADOS might perform as measures of symptomatology, especially if conducted within a restricted age range. Future work might also expand beyond cardinal diagnostic features and look into SC and RRB measured as quantitative autistic traits that expand beyond diagnostic features.

In conclusion, we have shown that SC-RRB balance can point to different macroscale functional connectivity phenotypes and potentially different genomic mechanisms that may underpin such phenotypes. While the divisions between these subtypes at the phenotypic level are not dramatically evident as categorical differences, at the level of macroscale neural circuitry, there is evidence to suggest that these SC-RRB subtypes are different when compared to the TD population. Future work to study these fractionable subtypes in an a-priori fashion will benefit from the use of our simple and supervised subtyping model and will further facilitate our understanding of how heterogeneity in autism manifests in a multi-scale fashion from genome to phenome.

## Acknowledgments

We thank all participants and their families for participating in this study. We also acknowledge the contributions of all members of the EU-AIMS LEAP group. Members of the EU-AIMS LEAP group are as follows: Jumana Ahmad, Sara Ambrosino, Bonnie Auyeung, Tobias Banaschewski, Simon Baron-Cohen, Sarah Baumeister, Christian F. Beckmann, Sven Bölte, Thomas Bourgeron, Carsten Bours, Michael Brammer, Daniel Brandeis, Claudia Brogna, Yvette de Bruijn, Jan K. Buitelaar, Bhismadev Chakrabarti, Tony Charman, Ineke Cornelissen, Daisy Crawley, Flavio Dell’Acqua, Guillaume Dumas, Sarah Durston, Christine Ecker, Jessica Faulkner, Vincent Frouin, Pilar Garcés, David Goyard, Lindsay Ham, Hannah Hayward, Joerg Hipp, Rosemary Holt, Mark H. Johnson, Emily J. H. Jones, Prantik Kundu, Meng-Chuan Lai, Xavier Liogier D’ardhuy, Michael V. Lombardo, Eva Loth, David J. Lythgoe, René Mandl, Andre Marquand, Luke Mason, Maarten Mennes, Andreas Meyer-Lindenberg, Carolin Moessnang, Nico Mueller, Declan G. M. Murphy, Bethany Oakley, Laurence O’Dwyer, Marianne Oldehinkel, Bob Oranje, Gahan Pandina, Antonio M. Persico, Barbara Ruggeri, Amber N. V. Ruigrok, Jessica Sabet, Roberto Sacco, Antonia San José Cáceres, Emily Simonoff, Will Spooren, Julian Tillmann, Roberto Toro, Heike Tost, Jack Waldman, Steve C. R. Williams, Caroline Wooldridge, and Marcel P. Zwiers.

## Supplementary Materials

**Fig. S1:**
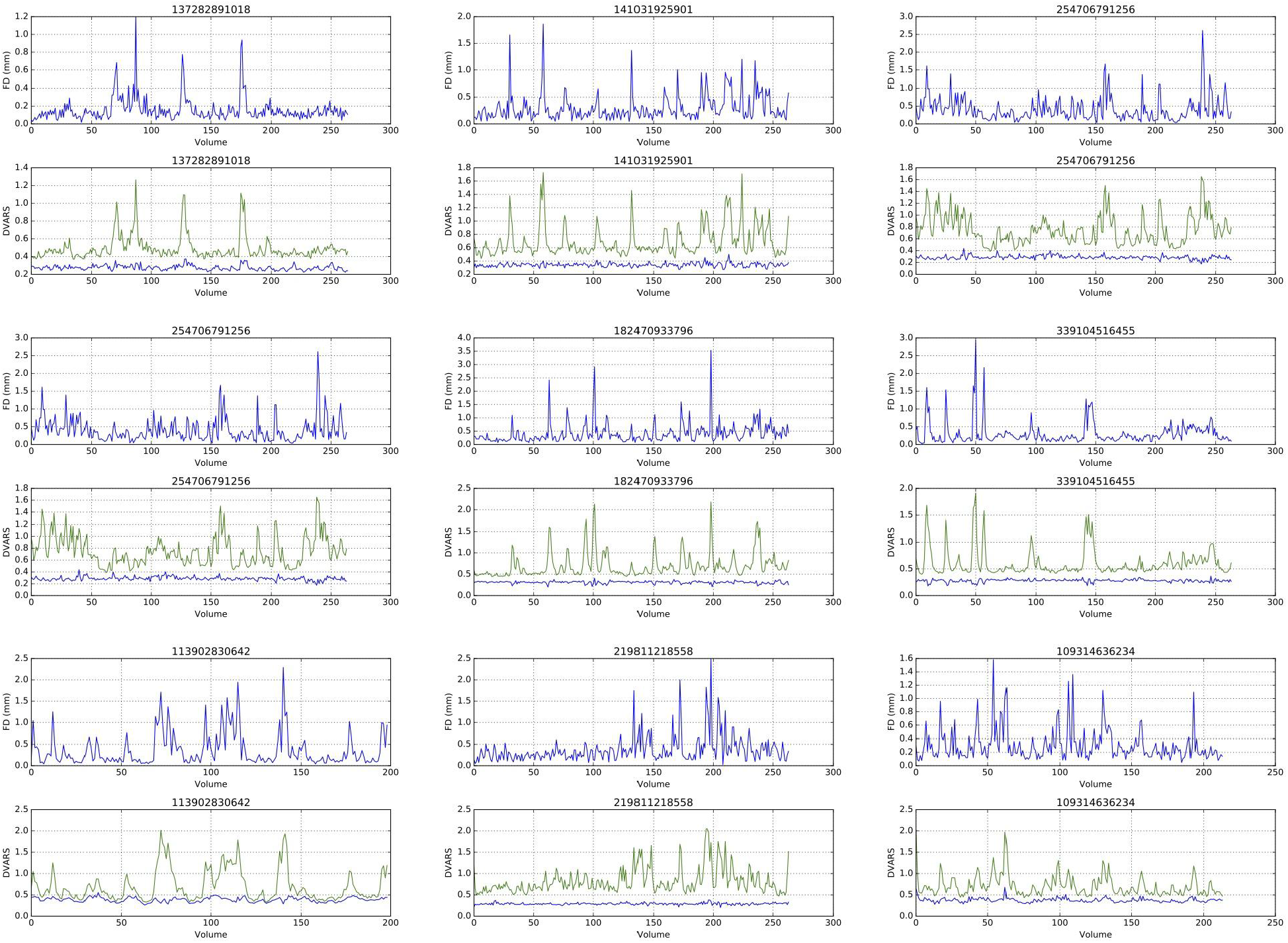
Examples of head-motion derived artefact that the effect of ME-ICA at flattening DVARS. This figure shows 9 example subjects, each with two plots. The top plot always shows framewise displacement (in mm) as an indicator of how much head motion is exhibited at each successive volume. The bottom plot shows DVARS traces from the optimally combined time-series before ME-ICA denoising (green) and after ME-ICA denoising (blue). DVARS traces before ME-ICA (green) closely follow the same pattern of framewise displacement and showcases how head motion can induce non-BOLD artefact. However, after ME-ICA denoising these DVARS traces are heavily flattened out, as a large proportion of this non-BOLD head motion artefact is isolated and removed as part of the denoising process.

**Fig. S2:**
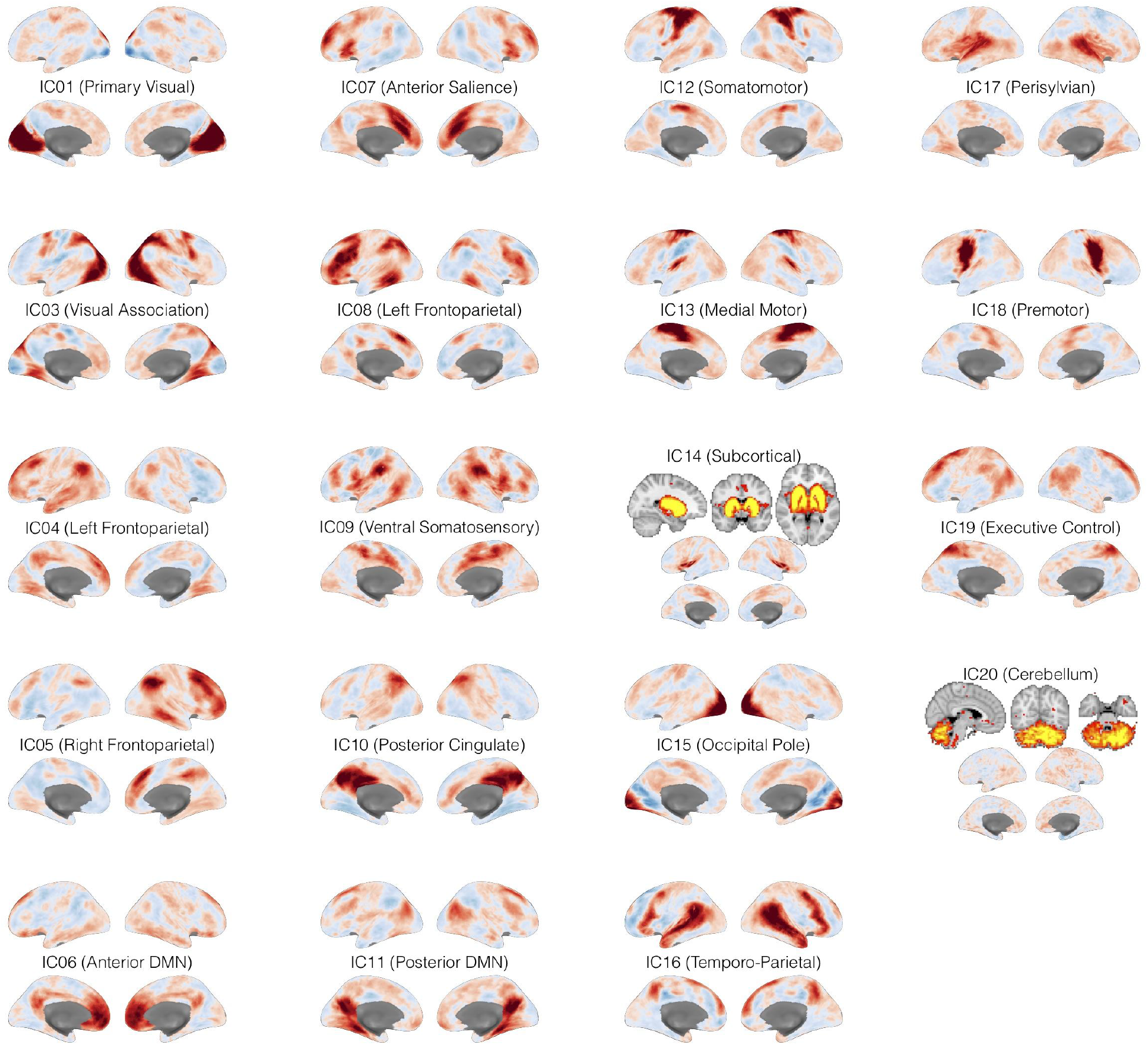
Visual depiction of the 19 components identified with group-ICA. Z-statistics are indicated with the blue-to-red color scale. Increasingly dark red regions are the regions of primary importance for the component. For components with mostly subcortical or cerebellar regions of importance (i.e. IC14 and IC20), these regions are highlighted in bright orange in axial, sagittal, and coronal planes.

**Fig. S3:**
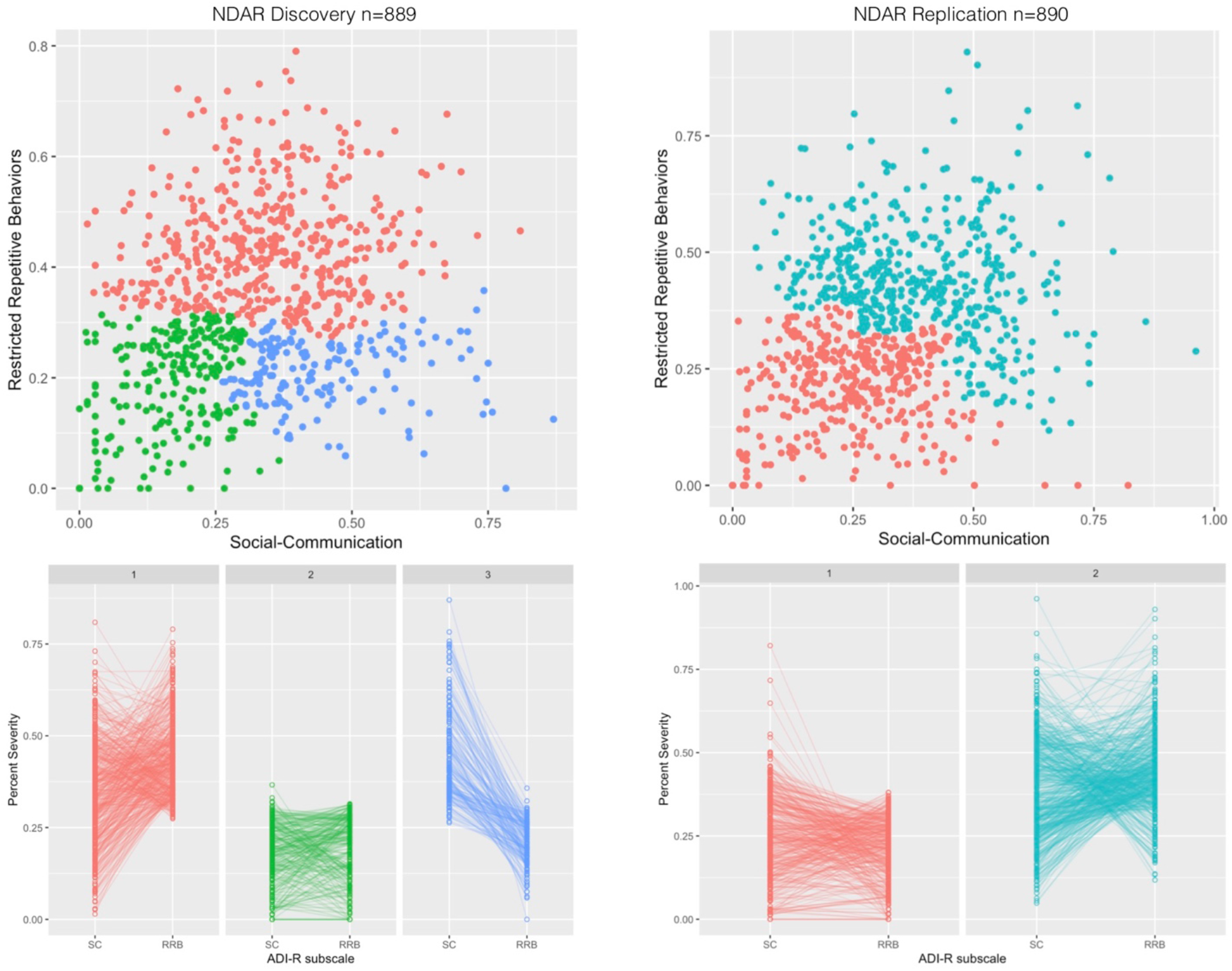
Unsupervised stratification with agglomerative hierarchical clustering with static cut height and internal clustering metrics to determine optimal number of clusters. This figure shows clustering results from NDAR Discovery and Replication datasets when the optimal number of clusters is selected by majority vote with NbClust.

**Fig. S4:**
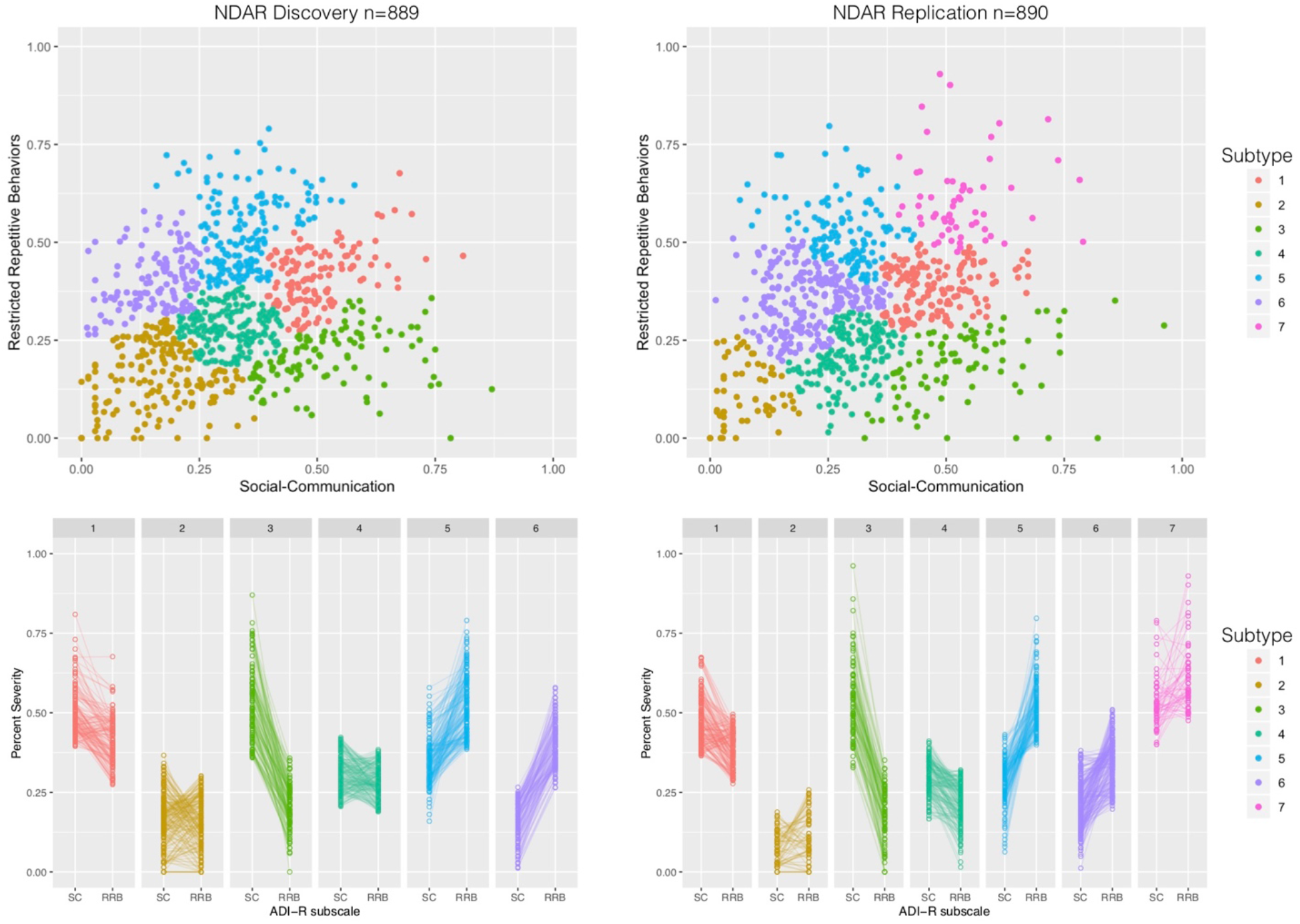
Unsupervised stratification with agglomerative hierarchical clustering and the dynamic hybrid tree cut algorithm for identifying clusters. This figure shows subtypes that emerge from clustering when the number of clusters is automatically selected with a dynamic hybrid tree cutting algorithm.

**Table S1:**
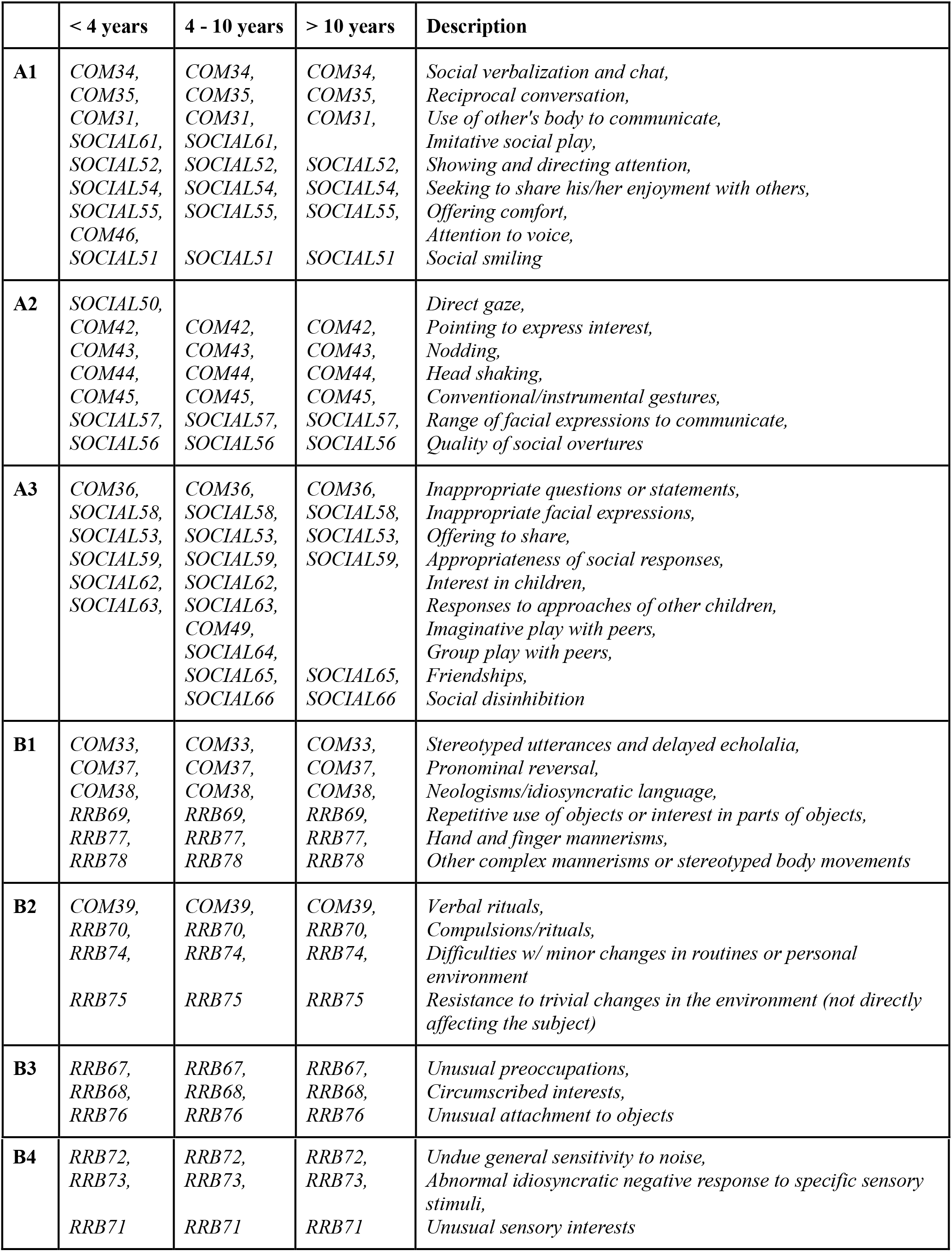
ADI-R items to use in DSM-5 scoring. This scoring scheme is identical to that reported by Huerta et al.,^26^. Subscales A1-A3 are within the social-communication (SC) domain while subscales B1-B4 are within the restricted repetitive behavior (RRB) domain. All items in A1-A3 utilize the Current scores, while all items in B1-B4 utilize the Ever scores.

**Table S2: Statistics from all functional connectivity comparisons.**

**Table S3: Gene lists used for enrichment analyses.**

